# Time-series analyses of directional sequence changes in SARS-CoV-2 genomes and an efficient search method for advantageous mutations for growth in human cells

**DOI:** 10.1101/2020.06.16.151282

**Authors:** Kennosuke Wada, Yoshiko Wada, Toshimichi Ikemura

## Abstract

We first conducted time-series analysis of mono- and dinucleotide composition for over 10,000 SARS-CoV-2 genomes, as well as over 1500 Zaire ebolavirus genomes, and found clear time-series changes in the compositions on a monthly basis, which should reflect viral adaptations for efficient growth in human cells. We next developed a sequence alignment free method that extensively searches for advantageous mutations and rank them in an increase level for their intrapopulation frequency. Time-series analysis of occurrences of oligonucleotides of diverse lengths for SARS-CoV-2 genomes revealed seven distinctive mutations that rapidly expanded their intrapopulation frequency and are thought to be candidates of advantageous mutations for the efficient growth in human cells.

## 1. Introduction

To confront the global threat of COVID-19 (World Health Organization, 2020), many SARS-CoV-2 genome sequences have been rapidly decoded and promptly released from the GISAI database (Elbe and Buckland-Merrett, 2017; Shu and McCauley, 2017). To elucidate characteristics of this virus in various aspects, we must introduce various research methods other than the phylogenetic tree construction, such as those used in big data analyses. Many host factors (e.g., nucleotide pools, proteins and RNAs) are involved in viral growth, and human cells may not present an ideal growth condition for invading viruses from nonhuman hosts. Therefore, efficient growth after the invasion will likely require changes in the virus genome. To study this viral adaptation, we previously analyzed time-series changes in mono- and oligonucleotide compositions of three zoonotic RNA viruses (Zaire ebolavirus, MERS coronavirus and influenza virus) and identified time-series directional changes that were detectable even on a monthly basis (Iwasaki et al., 2011, 2013; Wada et al., 2016). Additionally, an analysis of 25,000 influenza virus genomes showed that the directional changes were very similar between H1N1 (starting in 1918), H3N2 (starting in 1968) and H1N1/2009 (starting in 2009) (Wada et al., 2016). These reproducible changes of the three independent invasions should reflect viral adaptive processes for efficient growth in human cells.

## 2. Materials and methods

Genome sequences of human SARS-CoV-2 were downloaded from the GISAI database (https://www.gisaid.org/epiflu-applications/next-hcov-19-app/) (Elbe and Buckland-Merrett, 2017; Shu and McCauley, 2017); sequences belonging to the complete genome, high-coverage and human categories were downloaded on May 2, 2020. The polyA-tail sequence was removed before k-mer analyses. The Genome sequences of Zaire ebolavirus were downloaded from the Virus Variation Resource database (https://www.ncbi.nlm.nih.gov/genome/viruses/variation/) (Hatcher et al., 2017) on May 3, 2020.

In the study of RNA viruses, k-mers are subsequences of length k contained within the RNA virus genome. For an example of a very short sequence, the RNA sequence AGAUU has five monomers (A, G, A and 2Us), four 2-mers (AG, GA, AU and UU), three 3-mers (AGA, GAU and AUU), two 4-mers (AGAU and GAUU) and one 5-mer (AGAUU). For the k-mer counting of long genome sequences, KMC (http://sun.aei.polsl.pl/kmc) is a disk-based program for counting k-mers from FASTA files. Histogram analysis was done using the Excel function.

## 3. Results

### 3.1. Time-series changes in mononucleotide composition

To investigate whether time-series changes, which were previously found for Zaire ebolavirus, MERS coronavirus and influenza virus (Wada et al., 2016), occur also in SARS-CoV-2 isolated from humans, we plotted the mononucleotide composition (%) of each strain of SARS-CoV-2 in an order corresponding the isolation date, after the removal of the polyA-tail in advance (Fig. 1a). We first noticed the presence of the data that appear to be outliers and found that these were often derived from the sequences where the number of Ns (unknown nucleotides) is greater than 1000; this should be an unavoidable situation in the case of urgent sequencing, such as SARS-CoV-2. Even in the presence of the outliers, we have observed a time-series decrease in C% and an increase in U%, although slight.

**Fig. 1.**
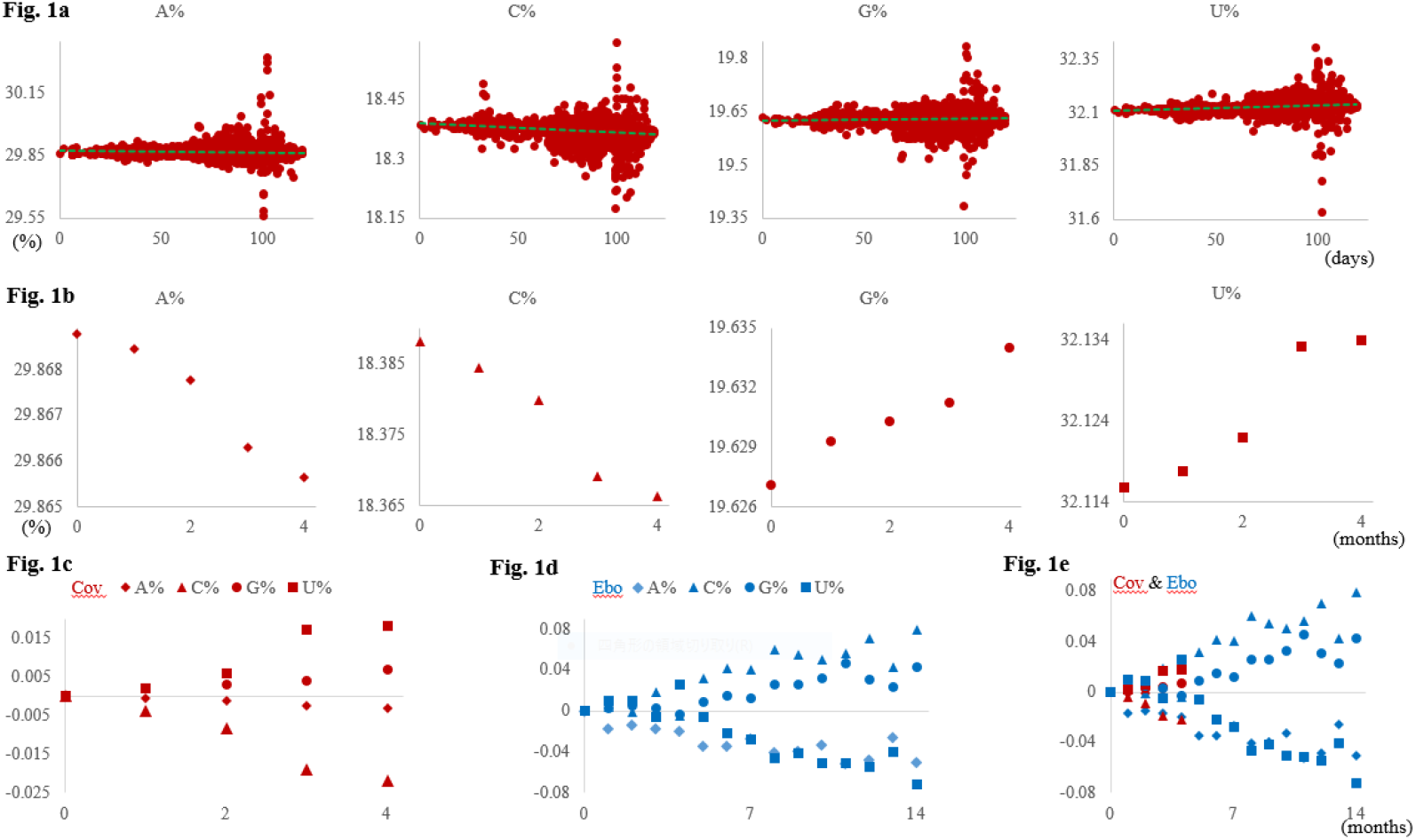
Time-series changes in the mononucleotide composition. (a) Mononucleotide composition (%) for each strain is plotted according to the days elapsed since 26/12/2019, the first day of SARS-CoV-2 isolation from human registered in the GISAI database. The regression line is indicated by a dashed line. (b) The averaged mononucleotide compositions (%) of SARS-CoV-2 strains tabulated for each month are plotted according to the elapsed months since December 2019; the 0 on the horizontal axis corresponds to December 2019; the 2 corresponds to February 2020; the 4 corresponds to April 2020. (c) The level of increase or decrease of each mononucleotide composition (%) of SARS-CoV-2 (Cov) from the start is plotted for each month. The colored symbols used for each mononucleotide are common with those used in b. (d) The increase or decrease level of each mononucleotide composition (%) of Zaire ebolavirus (Ebo) from its epidemic start (March 2014) is plotted for each month. The symbol shape is the same as SARS-CoV-2, but the color is changed to blue. (e) The increase or decrease level of each mononucleotide composition (%) from the start for the two viruses is plotted for each month. The colored symbols are common with those in c and d.

In the next analysis (Fig. 1b), rather than removing the noisy outliers, we tabulated the virus strains for each month of collection and calculated the mononucleotide composition (%) therein, as we previously done in the time series analysis of Zaire ebolavirus (Wada et al., 2016). The monthly results clearly show a monotonic decrease in A and C% and a monotonic increase in G and U%. This representation, however, does not directly compare a level of the change between mononucleotides. Using December 2019 as the starting point for the current pandemic (0 in both the horizontal and vertical axes), the level of increase or decrease of each mononucleotide composition (%) from the start is plotted for each month (Fig. 1c); the significant increase in G and U% and the decrease in A and C% can be visualized in a single figure.

Next, we compared the change observed for SARS-CoV-2 with that for Zaire ebolavirus, which started the epidemic in West Africa in March 2014. Although the number of ebolavirus sequences is much smaller than SARS-CoV-2, there are 15 months in which more than 10 strains per month have been reported, allowing us to see how long directional increases and decreases will be sustained. Using March 2014 as the start (0 in both the horizontal and vertical axes), the level of increase or decrease of each mononucleotide is plotted for each month (Fig. 1d). The previous study analyzed the three regions (Guinea, Liberia and Sierra Leone) separately and observed some regional differences, but the present study did not distinguish the three regions. Although there is some variation in data points that may be due to the regional difference and the small number of sequences, each nucleotide clearly undergoes a directional change (Fig. 1d). Fig. 1e shows both SARS-CoV-2 (reddish brown symbols) and ebolavirus (blue symbols) on a single figure. Since the shape of the symbol is identical in both viruses, it is clear that the direction of change in C% is reversed between the two viruses.

### 3.2. Time-series changes in dinucleotide composition

Next, we analyze the time series change in each dinucleotide composition (%) in the monthly tabulated SARS-CoV-2 and ebolavirus genomes. The first and second panels in Fig. 2a show the differences from the start month for 16 dinucleotides for SARS-CoV-2 and ebolavirus, respectively. In both viruses, various dinucleotides are clearly increased or decreased. Since the same symbol is used for the two viruses, it is clear that the direction of increase or decrease of individual dinucleotides differs between the two viruses. The last panel shows the data for both viruses, but here we have selected the first and second highest dinucleotides for the increase or decrease level in the last month of each virus. Although there are some differences between the dinucleotides, the trend of increase and decrease appears to slightly slow down after about six months from the ebolavirus epidemic start.

**Fig. 2.**
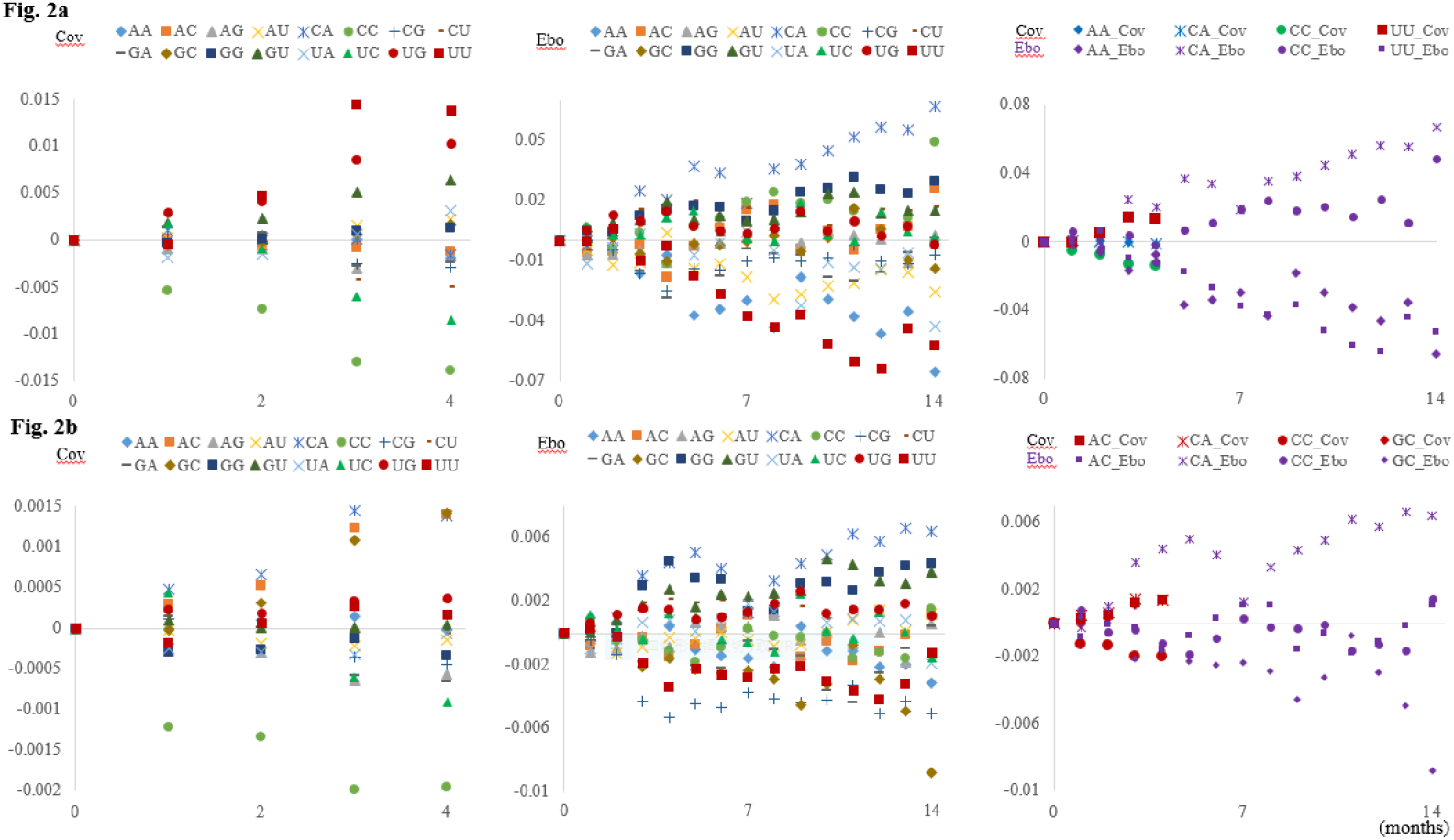
Time-series changes in the dinucleotide composition (%). (a) The increase or decrease level of each dinucleotide composition (%) from the start for the two viruses (Cov and Ebo) is plotted for each month. The last panel shows the data for both viruses; here we selected the first and second highest dinucleotides for the increase or decrease level in the last month of each virus; for ebolavirus, the symbols are miniaturized in violet to distinguish the data of two viruses. (b) The increase or decrease level of each month’s Obs/Exp value for each dinucleotide is plotted for the two viruses. The last panel shows the data for both viruses, as described in a.

Since the mononucleotide composition changes time-dependently (Fig. 1), it is obvious that the dinucleotide composition changes. However, there may exist some properties that become apparent only when analyzing continuous nucleotides. By focusing on the value (Obs/Exp) obtained by dividing the observed value of the dinucleotide (Obs) by the expected value from the mononucleotide composition (Exp), the effect of the mononucleotide change can be reduced. Fig. 2b shows the time-series difference in each month’s Obs/Exp value for each dinucleotide. In both viruses, there is a clear increase or decrease in various dinucleotides, but the types of dinucleotides that increase or decrease often differ from those observed in Fig. 2a and between the viruses. The last panel shows the data for both viruses; here we have selected again the first and second highest examples for the increase or decrease level. The time series pattern of Obs/Exp for ebolavirus in Fig. 2b appears to be more complex than that in Fig. 2a. When it comes to continuous nucleotides, the relationship to biological functions that include various molecular mechanisms of the viral adaptation to human cells should become more relevant than with mononucleotides, and this may be a reason for the complex pattern in Fig. 2b.

### 3.3. Rapidly increasing 20-mers

Sequential extension of the oligonucleotide length from dinucleotide will provide clues to elucidate the molecular adaptation mechanisms, but we next analyzed the occurrence of 20-mers in SARS-CoV-2 genomes, as previously conducted for ebolavirus and influenza virus (Wada et al., 2016, 2017). For these highly mutable viruses, the time-series analysis of long oligonucleotides such as 20-mers is unambiguously important for designing the PCR primers and therapeutic oligonucleotides that can be used for a sufficiently long period (Wada et al., 2017). Since the polyA-tails were removed in advance, none of the 20-mers was present more than once in each genome. Therefore, it is important to mention that the occurrence level (%) of a certain 20-mer in each month’s viral population corresponded to the occurrence level (%) of the strains with the 20-mer sequence in the viral population.

When we happened to search for 20-mers that were absent in all 16 strains isolated in December 2019, then emerged and increased in occurrence, we unexpectedly found a group of rapidly increasing 20-mers. This remarkable peculiarity is shown in Fig. 3a; here, we focused on all 39,990 types of 20-mers that were present in April 2020 but absent in December 2019 and analyzed their occurrence level (%) in the April population. The first histogram (reddish brown) displays the number of types of 20-mers present within each 3% width of the occurrence level in the April population. In the area with a very low occurrence level, the vertical axis value is near 40,000, which is almost the same to all 39,990 types emerged after December 2019; i.e., almost all 20-mers that newly emerged exhibit a very low occurrence level in the April population. However, when the vertical axis is displayed as a log (blue) scale, peculiar data (marked with a long arrow) become evident at the position where the horizontal value is more than 80% in the April population and the vertical log value is 1.903 (= 80). To be more precise, sixty types of 20-mers have an occurrence level of 82%, and twenty types have an occurrence level of 81%. When analyzing time-series changes in the occurrence level of the eighty 20-mers, a very similar pattern of monotonic increase was observed for all eighty 20-mers (Fig. 3b).

**Fig. 3.**
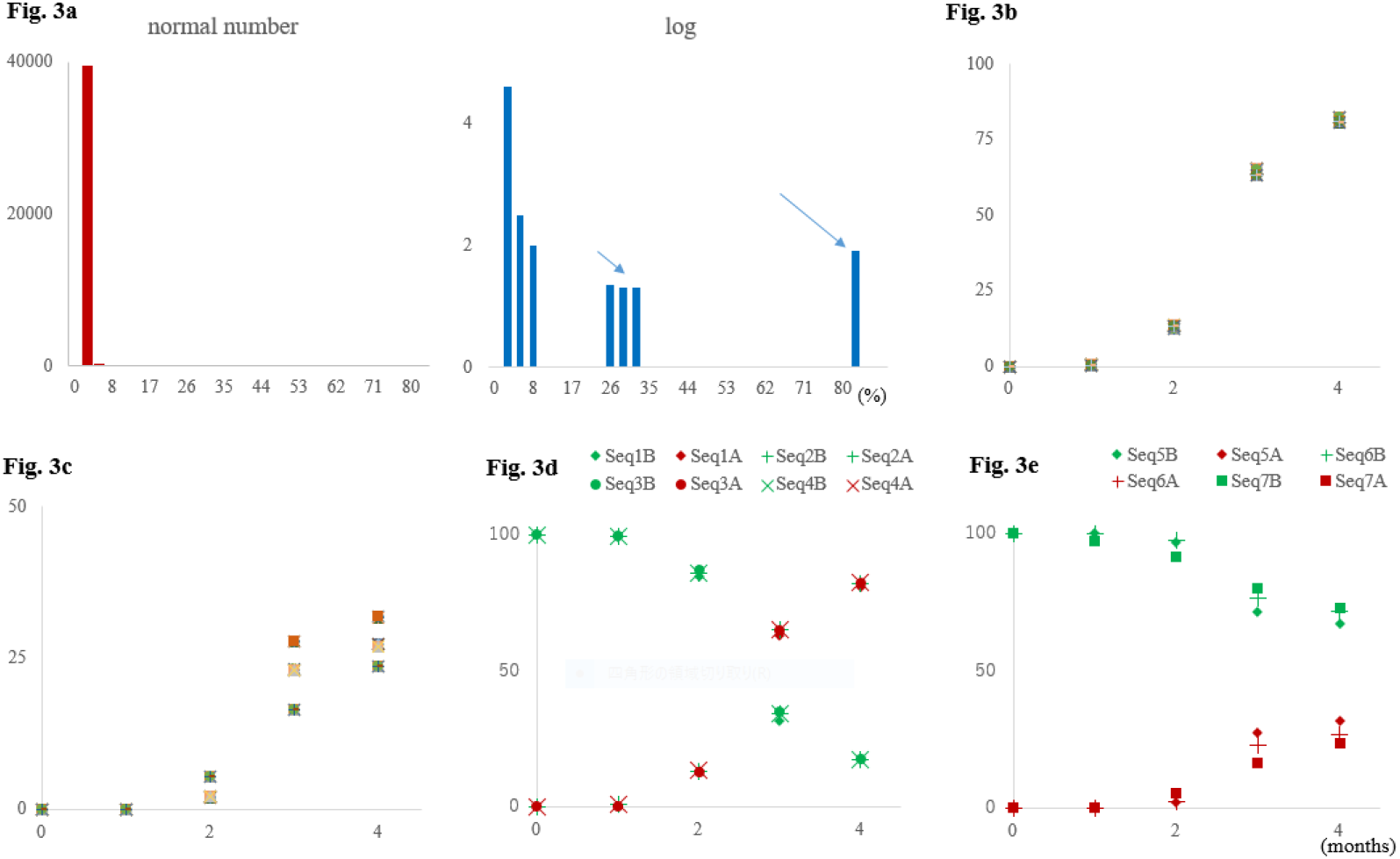
Histogram of the occurrence levels of 20-mers for April strains of SARS-CoV-2 and time-series changes of rapidly increasing 20-mers. (a) The vertical axis of the histogram shows normal numbers (reddish brown) or logarithms (blue); here, nonexistence in the logarithmic display is shown expediently as 0. Long and short arrows indicate the data locating in the highest and next highest region of interest. (b) Monthly occurrence of eighty 20-mers found in the highest region are plotted according to elapsed months. Since there is little difference between the eighty 20-mers, the relationship between each 20-mer and the symbol was omitted. (c) Monthly occurrence of 20-mers found in the second highest region (marked by a short arrow) are plotted according to elapsed months. The first, second and third rank in the last month correspond to 20-mers derived from Seq5, 6 and 7, respectively. While there is a slight difference in the intrapopulation frequency of each rank, resulting in a slight bulge (noted as a micro diversity in the text), the relationship between each 20-mer and the symbol is omitted as in b. (d) Monthly occurrences of pre- and post-mutation sequences of Seq1-4 are plotted according to elapsed months, respectively; green and reddish brown symbols are used for the pre- and post-mutation sequences with suffixes B and A, respectively. (e) Monthly occurrences of pre- and post-mutation sequences of Seq5-7 are plotted as described in d.

### 3.4. Genomic positions of rapidly increasing 20-mers and their time-series occurrences

We next focus on the eighty 20-mers that have rapidly and monotonically increased their intra-population frequency. Homology searches (BLASTn) against the viral genome sequences showed that they can be grouped into four sets of twenty 20-mers locating apart on the viral genome; the twenty 20-mers at each genomic location are shifted by one base, resulting in a single 39-mer. Figure 4 presents sequences of four 39-mers found at the four genomic positions: Seq1 in 5’ UTR, Seq2 in Orf1a for nsp3, Seq3 in Orf1b for RNA-dependent RNA polymerase and Seq4 in S for surface glycoprotein. Figure 4 presents also four pre-mutation sequences found in December strains in the top row. Figure 3c shows the time-series changes in a total of eight 39-mers; the sequences before and after the mutation (symbols in green and reddish brown, respectively) were distinguished by the additional letters B and A: e.g., Seq1B and 1A. The 39-mers observed before or after mutation show a clear monotonic decrease or increase, respectively; i.e., no bottleneck-type frequency reduction is observed for the postmutation 39-mers.

**Fig. 4.**
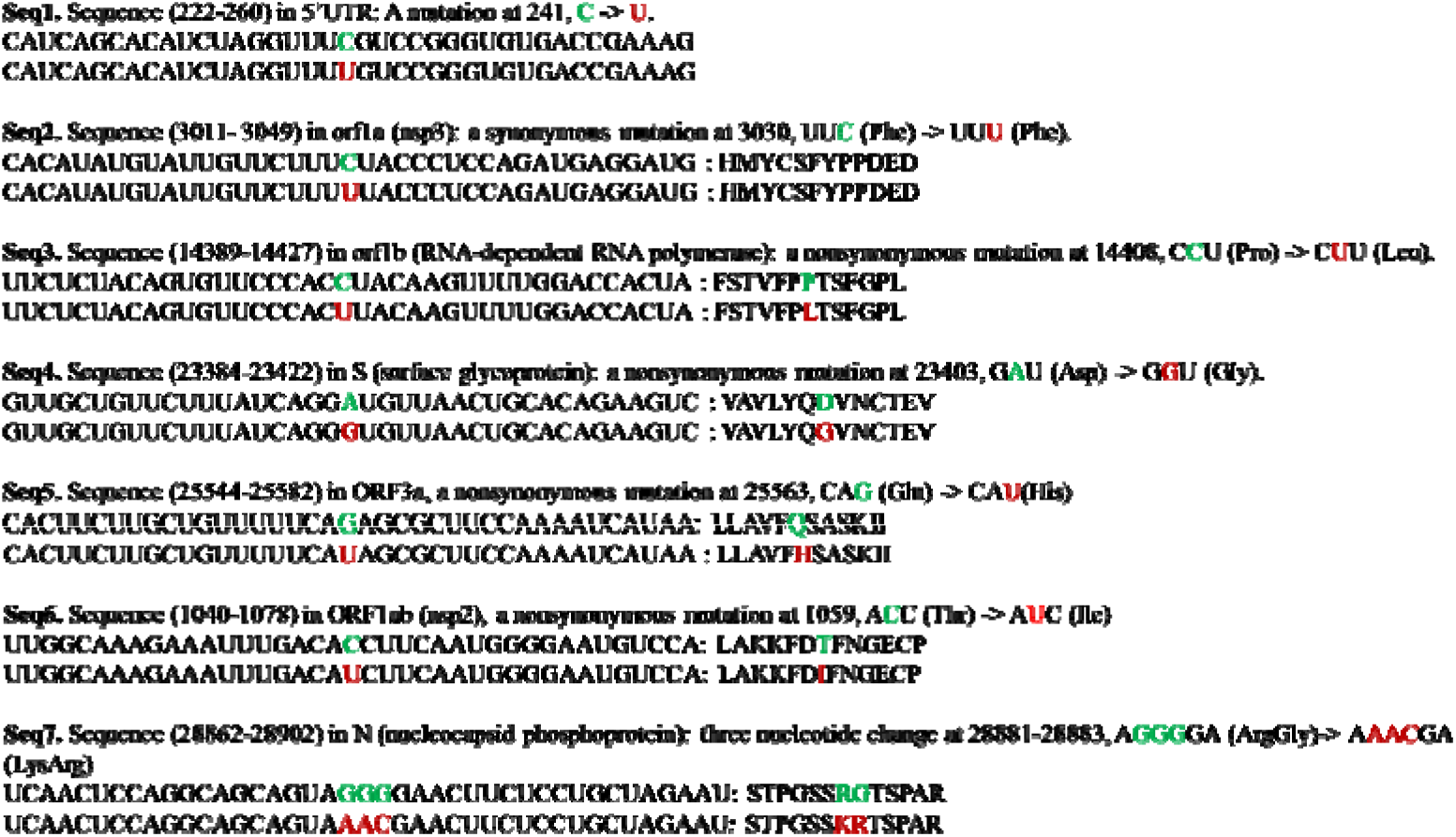
Sequences in focus and their genomic positions. The top row shows the premutation sequence found in December 2019 strains, and the bottom row shows the postmutation sequence found in April strains; the nucleotide in focus before or after the mutation is indicated in green or reddish brown. The nucleotide position was according to the reference genome (isolate Wuhan-Hu-1, EPI_ISL_402125) (Changtai, et al., 2020). The sequences other than the first sequence are from protein-coding genes, and the amino acid changes caused by the mutations are presented. Seq8 noted in Discussion was not detected in Fig. 3, because a portion of December strains had the sequence; the 29-mer sequence obtained in the 15-mer analysis is UAUGAGAACCAAAAAUUGAUUGCCAACCA: the sequence (24311-24339) in S encoding surface glycoprotein.

Looking at Fig. 3a, clearly distinctive data are also observed in the 32-24% range (marked with a short arrow), and the BLASTn analysis gave the results presented in Fig. 4. The twenty 20-mers with an occurrence level of 32% belonged to a 39-mers located in Orf3a (Seq5) and the twenty 20-mers with an occurrence level of 27% belonged to a 39-mers located in Orf1a for nsp2 (Seq6). In the case of the occurrence level of 24%, twenty-two 20-mers belonging to a 41-mers located in N for nucleocapsid phosphoprotein (Seq7). Interestingly, this mutation is not a point mutation, but three bases are changed at once, and therefore, the number of 20-mers with the mutation in focus increased from twenty to twenty two. Figure 3d shows the time-series changes in four 39-mers and two 41-mers listed in Fig. 4; those found before or after mutation show a clear monotonic decrease or increase, respectively.

Next, we re-examined the time series changes of the 20-mers (Fig. 3b and c) in more detail. Even for the 20-mers belonging to the same 39- or 41-mer, there is a slight difference in the occurrence level, which is observed as a slight bulge in the data at the same location in Fig. 3c. This slight diversity was found to be caused by mutations that occur around the interested mutations in a very small number of strains. Although the possibility of sequencing errors cannot be ruled out, we think this slight diversity reflects the random fluctuation caused by neutral mutations occurred around the interested mutations.

### 3.5. Search for the hidden advantageous mutations

To understand molecular mechanisms of viral adaptation to human cells, it is unambiguously important to find mutations that favor proliferation in human cells and human-to-human transmission. Suzuki and Gojobori (1999) developed a method for detecting positively selected sites in protein coding genes, by comparing the numbers of synonymous and nonsynonymous substitutions per site, and Suzuki (2006) applied it to influenza virus genomes. We next explain that the present histogram and time-series analysis, which does not rely on sequence alignment, is also a method for detecting positively selected sites in RNA virus genomes.

The most important characteristic of the positively selected advantageous mutations should be their rapid increase in the intra-population frequency. Seven mutations in focus are candidates for the advantageous mutations, but advantageous mutations are very rare compared to neutral mutations that are hitchhikingly synergistic to the driver-type advantageous mutation. It is thus important to distinguish the genuine advantageous mutation from the hitchhiker-type neutral mutations. To do so, we should thoroughly search for candidates for advantageous mutations and rank them in an order of the increase level of their intra-population frequency. If we can find such a mutation that increases the intra-population frequency more rapidly than the interested mutations, or at a similar level, the interested mutations in Fig. 4 may become the possible hitchhiker-type neutral mutations synergistic to the newly found mutation.

We first explain the characteristic of the present histogram analysis shown in Fig. 3a. There, we exhaustively analyzed all 20-mer sequences that were absent in all December (2019) strains but present in April (2020) strains. Therefore, the hidden advantageous mutation “X” is missed only when such mutable sites that can counteract the X’s intra-population expanse are present within 19 bases both upstream and downstream of the X mutation. To search for the X mutation, we should analyze the sequences shorter than 20-mer. Additionally, the restriction of non-existence in the December population is removed, because the X mutation may exist in a few December strains.

For all 15-mers found in SARS-CoV-2 genomes, Fig. 5a shows the histogram for their increase levels (%) in the April population from the December population. The pattern is very similar to that listed in Fig. 3a, and importantly, all cases of more than 20% increase are limited to the 107 (= 15×7 + 2) types of 15-mers with the seven mutations in focus; the addition of 2 in parentheses is due to a three-nucleotide change found in Seq7. It is worthy to note that ten out of the 59,922 type of 15-mers in April have two copies on the genome, but the two copies are also present in all December strains and their occurrence levels very slightly decrease in April, which does not affect the search for the X mutation. When it comes to 10-mers, more than 1200 out of a total of 47,002 types have multiple copies, but the 10-mers with the increase exceeding 20% are limited to 72 (= 10×7 + 2) with the interested mutations.

**Fig. 5.**
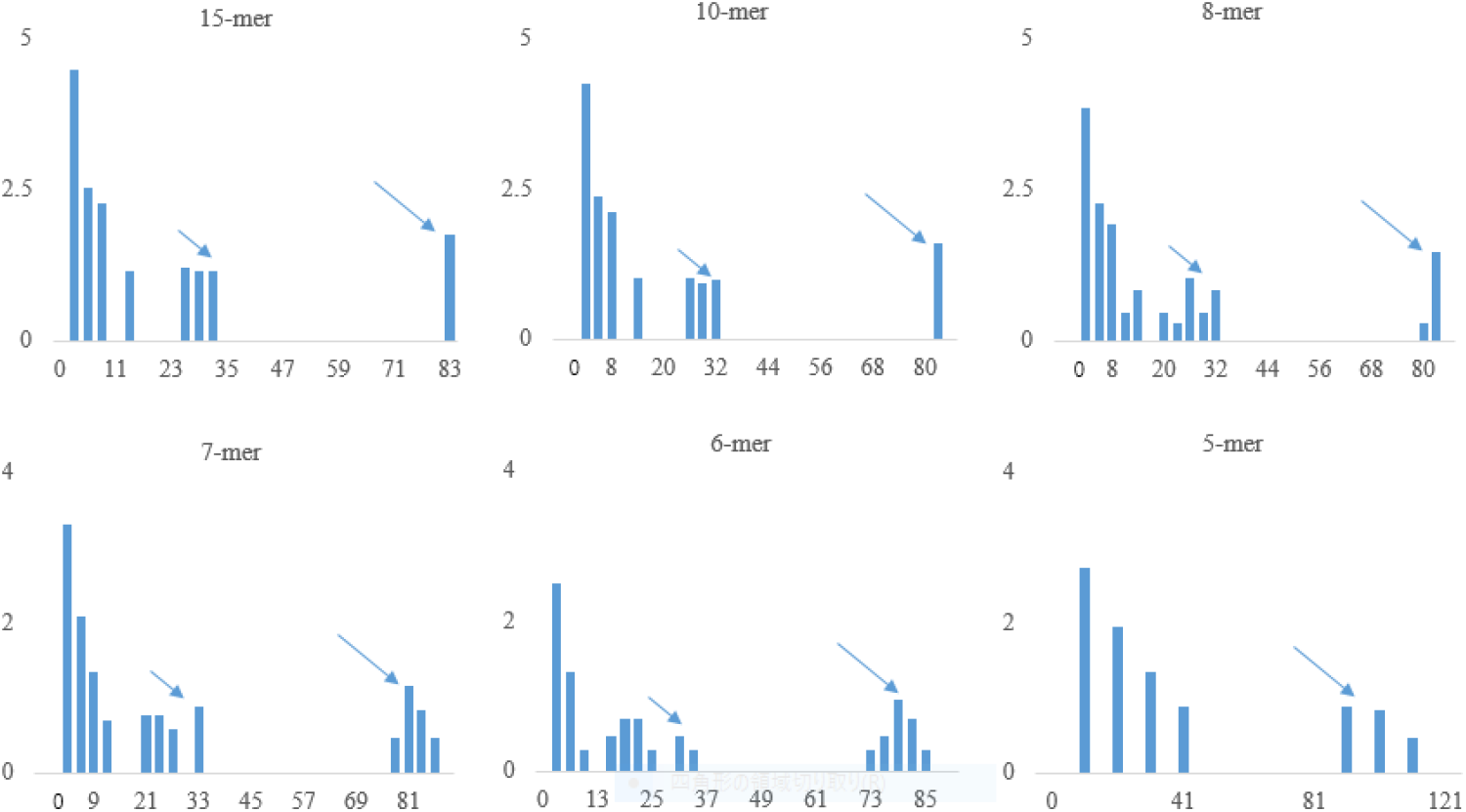
Histogram of occurrence levels of oligonucleotides shorter than 20-mer. The vertical axis of the histogram shows logarithms (blue), and nonexistence in the logarithmic display is shown expediently as 0. Long and short arrows indicate the data locating in the highest and next highest region of interest, as described in Fig. 3a.

In the case of 7-mer, approximately 60% of the 12,203 types present in April have multiple copies and some have more than twenty copies, and therefore, the result is expected to be complicated by the cumulative effect caused by many mutations occurring around the interested mutations. However, all 7-mers with more than 20% increase are limited to 51 (= 7×7 + 2) types with the interested mutations, showing the extraordinarily rapid increase in their intra-population frequency. Even though, the histogram pattern has changed significantly; the region of the high values (marked by a long arrow) has begun to split, with the highest value reaching near 90%. It should be noted that the top three were GGGUGUU, UUUUUAC and CUUUUUA (i.e., U- and/or G-rich sequences), whose increase is predicted from the mono- and dinucleotide increase (Figs. 1 and 2). In other words, the effect of the mono- and dinucleotide increase is also becoming apparent.

Most 5-mers are present in more than one hundred copies on the genome, and a few have ten thousand copies. Despite this, the top 20 (= 5×4) have the mutations in Seq1-4. However, the distribution in the histogram is continuous in the region below 40%, and the 5-mers with and without the mutations in Seq5-7 were mixed.

## 4. Discussion

### 4.1. Oligonucleotides shorter than 5-mer

When shortening the oligonucleotide length to 4-mer, there appeared two cases with over 200% increase, and this was because the highest UUUU has the mutation in both Seq1 and 2 and the second highest UUAC has the mutation in both Seq2 and 3 (Fig. 4). In addition, UGUU with no mutation in Seq1-4 appeared in the sixth position; UGUU is the most abundant 4-mer in April, with an average of over 300 copies per genome, and its increase is predictable from the mono- and dinucleotide increase. Even when shortening the oligonucleotide length to 3-mer, the top five had the Seq1-4 mutations. We thus concluded that there is no hidden advantageous mutation that expands its intra-population frequency at a faster rate than the Seq1-4 mutations, during the current SARS-CoV-2 pandemic to April 2020.

Next, we discuss the mutations in Seq5-7. When shortening the length to 4-mer, some 4-mers increased at the same level as those for the Seq5-7 mutations, and interestingly, such cases that were not expectable from the mono- and dinucleotide increase were observed in a high order: e.g., AAAU and AAUU. Interestingly, these corresponded to the 4-mers that constitute the 15-mers, whose increase level was in the eighth position (12% increase in Fig. 5); for detail of the Seq8, see the legend to Fig. 4. We thus concluded that there is no hidden advantageous mutation that expands its intra-population frequency at a faster rate than the mutations in Seq5-7.

### 4.2. Characteristic of sequences determined by the PCR method and order of appearance of the interested mutations

First, we should point out that the viral sequence determined by the PCR method is of an average property of multiple viral copies isolated from one patient. Even if a neutral mutation occurs in a certain viral genome, unless the copy number at the start of the infection is relatively low and the neutral mutation occurs early in the infection, the mutation is not reflected in the determined sequence of the patient because of the PCR averaging operation.

Next, we explain the order of the mutations found in Seq1-7. The mutations in Seq1-4 appeared first in the following two January strains; the hCoV-19/Germany/BavPat1/2020|EPI_ISL_406862|2020-01-28 carried mutations in Seq1, 2 and 4, and the hCoV-19/Zhejiang/HZ103/2020|EPI_ISL_422425|2020-01-24 carried mutations in Seq2, 3 and 4; i.e., mutations in Seq2 and 4 were commonly present in these two strains. However, the mutation in Seq5-7 had not yet emerged in January. Among 493 strains in February, mutations in Seq1, 2, 3 and 4 were found in 65, 65, 62 and 67 strains. New mutations in Seq5, 6 and 7 were found in 10, 11 and 27 strains, and were almost always present in strains with the Seq4 mutation; i.e., Seq5, 6 and 7 mutations occurred in a strain with the Seq4 mutation. Although the majority of Seq5 and 6 mutations were observed in common strains, the Seq7 mutation was exclusive to the former two; i.e., the mutation occurred independently from those of Seq5 and 6.

A more detailed discussion will require epidemiological information, but from the results of the time series analysis alone, we can conclude that the Seq4 mutation is the genuine advantageous mutation, most significantly increasing the intrapopulation frequency from January to April. The Seq7 mutation is also considered to be an advantageous mutation, because this mutation appeared after the strain with the Seq4 mutation has reached a significant number. While the possibility that either Seq5 or 6 is a neutral mutation cannot be ruled out, both are considered to be advantageous mutations because the mutations are nonsynonymous between amino acids with different chemical properties.

### 4.3. A sequence alignment-free method for searching advantageous mutations

As explained in Introduction, a wide variety of technologies must be introduced to combat emerging infectious diseases, such as COVD-19. The present study has attempted to develop a sequence alignment free method that will become a complement to the phylogenetic tree construction in viral evolutionary studies. We focused on the technological development, rather than discussing biological implications of the obtained results. Time-series analysis of occurrences of oligonucleotides of diverse lengths can unveil various characteristics hidden in the virus genome. Furthermore, the present method can extensively search for advantageous mutations (i.e., unveil the hidden ones) and rank them in an increase level for their intra-population frequency. All methods used here are suitable for big data analysis. In addition, although there are a significant number of viral sequences that contained many Ns, the present analysis does not require any special preprocessing. As a characteristic of big data analysis, the effects of erroneous data naturally decrease with the increase in the data size. The present method should become a powerful complement to the phylogenetic tree method in the viral evolutionary study. Combining it with an unsupervised-type AI (Iwasaki et al., 2011, 2013) can provide a powerful method for obtaining novel knowledge from big data of viral genome sequences.

## Acknowledgements

We gratefully acknowledge the authors submitting their sequences from GISAID’s Database. We gratefully acknowledge also the valuable comments of Dr. Yashushi Hiromi of National Institute of Genetics (Mishima). This work was supported by a Grant-in-Aid for Scientific Research (18K07151) from the Ministry of Education, Culture, Sports, Science and Technology, Japan.

## References

Elbe S., Buckland-Merrett G., 2017. Data, disease and diplomacy: GISAID’s innovative contribution to global health. Global Challenges, 1, 33–46. doi:10.1002/gch2.1018 PMCID: 31565258.

Hatcher E.L., Zhdanov S.A., Bao Y., Blinkova O., Nawrocki E.P., Ostapchuck Y., Schäffer A.A., Brister J.R., 2017. Virus Variation Resource - improved response to emergent viral outbreaks. Nucl. Acids Res. 45(D1), D482–D490. doi: 10.1093/nar/gkw1065.

Iwasaki Y., Abe T., Wada K., Itoh M., Ikemura T., 2011. Prediction of directional changes of influenza A virus genome sequences with emphasis on pandemic H1N1/09 as a model case. DNA Res. 18, 125–136.

Iwasaki Y., Abe T., Wada Y., Wada K., Ikemura T., 2013. Novel bioinformatics strategies for prediction of directional sequence changes in influenza virus genomes and for surveillance of potentially hazardous strains. BMC Infect. Dis. 13, 386.

Shu Y., McCauley J., 2017. GISAID: Global initiative on sharing all influenza data – from vision to reality. EuroSurveillance, 22(13) doi: 10.2807/1560-7917.ES.2017.22.13.30494 PMCID: PMC5388101

Suzuki Y., 2006. Natural selection on the influenza virus genome. Mol. Biol. Evol. 23, 1902–1911.

Suzuki Y., Gojobori T. 1999. A method for detecting positive selection at single amino acid sites. Mol. Biol. Evol. 16, 1315–1328.

Wada Y., Wada K., Iwasaki Y., Kanaya S., Ikemura T., 2016. Directional and reoccurring sequence change in zoonotic RNA virus genomes visualized by time-series word count. Sci. Rep. 6, 36197.

Wada K., Wada Y., Iwasaki Y., Ikemura T., 2017. Time-series oligonucleotide count to assign antiviral siRNAs with long utility fit in the big data era. Gene Ther. 24, 668–673.

Wang C., Liu Z., Chen Z., Huang X., Xu M., He T., Zhang Z., 2020. The establishment of reference sequence for SARS-CoV-2 and variation analysis. J. Med. Virol. doi: 10.1002/jmv.25762.

World Health Organization. 2020. Coronavirus Disease (COVID-2019) Situation Report.

